# Single-cell transitional dynamics unravel stimulus- and cell type-dependent signaling outputs of distinct p53 regulatory feedback motifs

**DOI:** 10.1101/2021.02.08.430353

**Authors:** Jun Xie, Lichun Zhang, Yanting Zhu, Xiao Liang, Jue Shi

## Abstract

Despite myriad of systemic analysis, our quantitative understanding of signaling networks and their differential outputs is still limited, especially at the dynamic level. Here we employed a network motif approach to dissect key regulatory units of p53 pathway and determined how their collective activities generated context-specific dynamic responses. By combining single-cell imaging and mathematical modeling of dose-dependent p53 dynamics induced by three chemotherapeutics of distinct mechanism-of-action, including Etoposide, Nutlin-3a and 5-Fluouracil, and in five cancer cell types, we identified novel p53 dynamic modes and their mechanistic origins with new insight on previously unknown drug mediators and phenotypic heterogeneity of cancer cells. Our results established the functional roles of unique feedback motifs of p53 pathway in generating the stimulus- and cell type-specific signaling responses and demonstrated that transitional dynamics activated at intermediate stimulus levels can be exploited as novel quantitative readouts to uncover and elucidate the key building blocks of bio-networks.

## Introduction

The p53 regulatory pathway, consisting of the key tumor suppressor and transcription factor p53, its upstream regulators and downstream target genes, plays a central role in eliciting proper cellular responses to chromosomal damage and various stresses in mammalian cells [1, 2]. Referred to as the “guardian of the genome”, p53 and the extended p53 pathway have been widely studied, revealing multi-level post-transcriptional control over p53 activity via a complex network of signaling kinases, cofactors and inhibitory partners [3–5]. The plethora, and sometimes conflicting, data regarding p53 pathway-mediated signaling responses also prompted quantitative, systemic analysis by mathematical modeling and bioinformatics analysis to decipher the differential control of p53 pathway activity at the collective, network level [6–9]. However, most of the computational work employs the network topology-based approach and falls short of providing mechanistic insight, in particular for understanding the divergent p53 pathway responses to different stress stimuli and in variable mammalian cell types.

Single cell studies both by us and others showed that alteration of the induction dynamics of p53 not only is an important mechanism to modulate p53 activity but also provides a quantitative, dynamic readout to investigate the unique network motifs and regulatory modules of p53 pathway, alternative to the static network topology-based modeling approach. The control of p53 dynamics over cell fate was first examined for cellular response to transient gamma or UV irradiation and revealed intriguing periodic oscillation of p53 level [10–13]. We later uncovered other p53 dynamic modes in response to DNA damaging drugs, i.e., monotonic p53 induction and an extended large p53 pulse, which led to cell death and cell cycle arrest, respectively. We also found activation of the distinct p53 dynamic modes was context specific, depending on not only the DNA damage level but also the mammalian cell type [14, 15]. However, the studies of p53 dynamics so far are mostly on response to DNA damage, and we know much less about how p53 pathway and p53 dynamics are differentially activated by other types of stress stimuli.

On the mathematical modeling of p53 pathway dynamics, previous work largely focused on the periodic pulsing mode of p53, attributing it to a time-delay negative feedback loop between p53 and its main negative regulator and transcriptional target, Mdm2 [10, 16–18]. We found that an additional inhibitory interaction between ATM and Mdm2, resulted from ATM-mediated Mdm2 degradation, synergized with the p53-Mdm2 negative feedback to constitute a positive feed-forward motif in an ATM/p53/Mdm2 regulatory module and switched the pulsing mode to monotonic induction of p53 upon increasing DNA damage level [15]. However, while modeling and simulation of this regulatory module generated p53 dynamics largely similar to the single cell imaging results at low and high DNA damage level (i.e., the saturating damage dose regime), the model predicted a unique transitional dynamic phenotype of p53 at the intermediate DNA damage level, with an initial p53 pulse followed by an elevated plateau of p53 level, which we did not observe experimentally. For all the mammalian cell lines that we studied so far, the cell population shifted directly from p53 pulsing to monotonic induction that promotes cell death or an extended large pulse that promotes cell cycle arrest, without exhibiting a different mode of transitional dynamics [15]. This discrepancy led us to hypothesize that additional regulatory components/motifs beyond the well-known ATM/p53/Mdm2 regulatory module is involved in generating the DNA damage-induced bimodal switch of p53 dynamics. The unique transitional dynamics of p53 may also provide a new quantitative readout, in addition to the commonly studied dynamic characteristics, such as signaling noise, pulsing periods and amplitudes, to investigate a broader spectrum of p53 pathway motifs and their functional impacts.

We thus set out in this study to examine and elucidate the variable signaling outputs of p53 pathway in a broad cellular context, using three anticancer chemotherapeutics that activate p53 via distinct mechanisms, including Etoposide (DNA damage inducer), Nutlin-3a (Mdm2 inhibitor) and 5-Fluouracil (nucleolar stress inducer), and five human cancer cell lines. Our single cell imaging data revealed novel p53 dynamic modes that are stimulus- and cell type-dependent, in particular intriguing p53 transitional dynamics induced at intermediate drug concentrations. Importantly, it is these new p53 dynamic modes that vary significantly between drugs and cancer cell types, not the widely studied periodic pulsing mode of p53. Based on the experimental data, we then constructed and simulated the minimal, core regulatory modules of p53 pathway in a large parameter space, which allowed us to characterize unique feedback motifs and uncover their respective quantitative roles in generating the differential dose response of p53 dynamics in a context specific manner. Our results also revealed previously unknown mediators of the chemo-drug actions that vary between cancer cell types and impact drug sensitivity. For instance, we found MCF7, the cell line used in most of the previous studies of p53 response dynamics, is unique in its strong dependence on the negative feedback motifs, thus exhibiting resistance to drug-induced p53 upregulation and subsequent cell death in response to all three anticancer drugs, despite the different drug mechanisms. Our work provides important new insight into not only the context-specific p53 pathway control at the collective, modular level but also a quantitative framework to dissect the functional building blocks of cellular network via the dynamics of key signaling proteins.

## Results

### Stimulus- and cell type-dependent p53 dynamics

To experimentally profile the differential signaling outputs of p53 pathway, we used single-cell p53 dynamics acquired from live cell imaging as the quantitative signaling readout. We activated p53 and the p53 pathway by three different anticancer drugs. The DNA damaging drug, Etoposide, is a topoisomerase inhibitor and activates the DNA damage signaling kinase, ATM, which then phosphorylates both p53 and Mdm2, subsequently attenuating the inhibitory p53-Mdm2 interaction that enables the induction of p53 [19, 20] (Fig. 1A). The Mdm2 inhibitor, Nutlin-3a, directly abrogates the binding of Mdm2 to p53, thus allowing p53 level to increase [21, 22] (Fig. 1B). The anti-metabolite chemotherapeutic, 5-Fluouracile (5-FU), is a pyrimidine analog that interferes with DNA and RNA synthesis [23]. Although 5-FU induces both DNA and RNA damage, increasing evidence suggests that p53-mediated 5-FU cytotoxicity is mainly triggered by impairment of ribosomal biosynthesis in the nucleoli [24]. Such nucleolar stress activates the translocation of ribosomal proteins, such as RPL11 and RPL5, to bind and sequester Mdm2, leading to p53 upregulation [25–27] (Fig. 1C). In addition to the different drugs, we also characterized the cell-type variation in p53 pathway response by employing five p53 reporter cancer cell lines that we studied before, including A549 (lung), U-2 OS (bone), A375 (skin), MCF7 (breast) and 769-P (kidney) [15]. These cell lines were engineered to stably express a p53-Venus construct (i.e. wild-type p53 tagged with a yellow fluorescent protein, Venus) [11, 14]. Time-lapse imaging of the respective clonal fluorescent reporter lines allows us to obtain real-time p53 dynamics in individual cells via the fluorescent signal of p53-Venus, which is mostly localized in the nucleus. Representative still images and quantified time courses of the average nuclear p53-Venus fluorescence that the majority of cell population exhibited at each drug concentration are shown in Figure 1 (middle and right panels) for the selected three drugs and five cancer cell lines.

**Figure 1.**
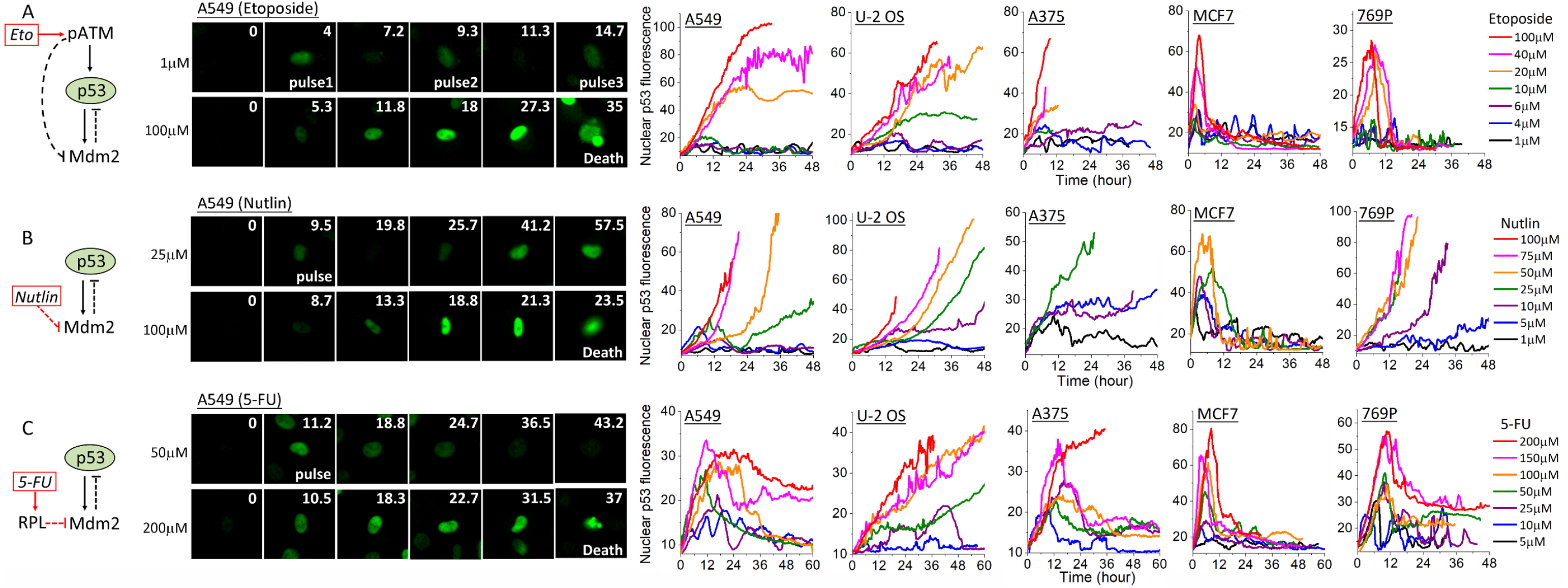
Dose responses of p53 dynamics to (A) Etoposide, (B) Nutlin-3a and (C) 5-Fluouracil (5-FU) vary between drugs and cancer cell types. <u>Left panel:</u> Pathway diagrams of key regulatory components of the p53 pathway known to mediate the respective drug response. <u>Middle panel:</u> Still cell images of p53 fluorescence monitored by the p53-Venue reporter in A549 cells at the indicated drug concentrations. Still images were cropped from time-lapse movies. Time (unit: hour) is indicated at the top right corner of each image. <u>Right Panel:</u> Representative single-cell trajectories of p53 dynamics in the nucleus quantified from p53-Venus fluorescence in the five cancer cell lines, including A549, U-2 OS, A375, MCF7 and 769-P. p53 dynamic responses to the different drug concentrations are color coded as shown on the figure. Cells were treated with the respective drug at time 0 and tracked for 48 to 60 hours or till cell death occurred. The abrupt end of some p53 trajectories before 48 hour, especially under high drug concentrations, corresponds to the time of death.

The single cell data revealed both stimulus- and cell type-dependent characteristics in the dose responses of p53 dynamics (Fig. 1). At low dose, Etoposide, Nutlin-3a and 5-FU all triggered periodic pulsing of p53 in all five cell lines, although some cell lines and drugs appeared to engender more regular periodic pulsing (e.g., A549 under 1 μM Etoposide). Upon increasing drug concentrations, p53 induction changed to different dynamic modes that varied between both the drugs and cell lines. Under Etoposide, the cell population switched to either monotonic induction of p53 (A549, U-2 OS and A375) or an extended large p53 pulse (MCF7 and 769-P), without exhibiting alternative transitional dynamics between the low and high drug concentrations (Fig. 1A, right panel). In contrast, responses to Nutlin-3a showed a distinctive transitional mode of p53 dynamics at the intermediate drug doses in A549, U-2 OS and 769-P cells before a further change to the saturating, high-drug mode. Specifically, for these three cell lines, intermediate concentrations of Nutlin-3a (10-50 μM) activated an initial p53 pulse followed by approximately exponential upregulation of p53, a novel p53 dynamic mode not observed before (Fig. 1B, right panel). As Nutlin-3a concentration further increased, the initial p53 pulse vanished and p53 induction kinetic was largely exponential. Dose responses to 5-FU revealed yet another new mode of transitional dynamics in A549, A375 and 769-P, in which the initial p53 pulse was followed by an elevated plateau of p53 level that was lower than the amplitude of the initial pulse (Fig. 1C).

In addition to drug-dependent variability, significant cell-type variation in p53 responses was also observed between the five cell lines in response to the same drug that correlated with alternative cell fate outcome of cell cycle arrest and cell death. p53 dynamics induced by Etoposide exhibited two distinct dose-response phenotypes that varied between cell types, one was from p53 pulsing to monotonic induction (A549, U-2 OS, A375) and the other was from pulsing to an extended large pulse (MCF7 and 769-P). We have previously shown that the monotonic induction mode, which resulted in high level of p53 upregulation, promoted drug-induced cell death, while the extended large pulse mode that restrained p53 upregulation at a low level led to cell cycle arrest, rendering MCF7 and 769-P resistant to Etoposide-induced cell death even at high drug concentrations [15]. Treatment of Nutlin-3a and 5-FU resulted in three different dose-response phenotypes of p53, including the unique transitional dynamic modes as discussed above and the similar saturating dynamic modes as seen in Etoposide. Interestingly, MCF7 cells showed largely the same dose-dependent p53 dynamics in response to all three drugs, despite the different mechanisms of action of these drugs, suggesting that MCF7 may have evolved to regulate p53 upregulation by one dominant regulatory motif. Moreover, MCF7 exhibited the suppressive extended large pulse mode of p53 dynamics at saturating high concentrations of all three drugs, which led to mostly cell cycle arrest and resistance to drug-induced cell death. In contrast, A549 and 769-P cells showed three distinct dose responses of p53 dynamics to the three different drugs, indicating their functional dependence on a more diverse set of p53 regulatory motifs, which could be potentially exploited, e.g., to confer drug selectivity. For instance, 769-P was resistant to Etoposide-induced cell death due to the suppressive, low p53 upregulation in the form of extended large pulse. In contrast, p53 in 769-P can be upregulated to a high-level, monotonic induction mode by high doses of Nutlin-3a, followed by cell death (indicated by the abrupt end of the p53 trajectories before 36 hours in Fig. 1), illustrating that Nutlin-3a is a more effective cytotoxic therapeutic for 769-P. Overall, the striking variability in the dose responses of p53 dynamics exhibited by the different cell lines suggested that p53 activation is regulated not only by variable drug mechanisms but also the cellular contexts of the mammalian cells, which likely have evolved differential dependence on selected p53 pathway components and interaction motifs. As the variable p53 levels resulted from the distinct p53 dynamics played a key role in promoting alternative cell fate outcome of cell cycle arrest and cell death, as we elucidated before [15], it is important to understand the quantitative and mechanistic basis of the differential dynamic outputs of p53 pathway, which we elaborated below.

### Dynamic output of the core p53-Mdm2 negative feedback loop

To elucidate the mechanistic origin of the stimulus- and cell type-dependent dose response of p53 dynamics that we experimentally observed at the network motif level, we first analyzed the dynamic output of the core p53-Mdm2 negative feedback loop. Etoposide perturbs this negative feedback via phosphorylation and activation of ATM, while Nutlin-3a and 5-FU inhibit Mdm2 directly (Fig. 1, left panel). We performed computational analysis of both regulatory motifs, and found their quantitative outputs of p53 dynamics were largely similar. As we already detailed the mathematical formulation of the ATM/p53/Mdm2 module in a previous study [15], here we elaborated the kinetic equations and simulation results in response to Nutlin-3a as a representative scenario. Results for the p53-Mdm2 response to Etoposide by activating the ATM/p53/Mdm2 module are provided in the supplementary materials (suppl. Fig.S1).

Dynamic output of the p53-Mdm2 negative feedback in response to Nutlin-3a is formulated by the following delay differential equations (DDEs).

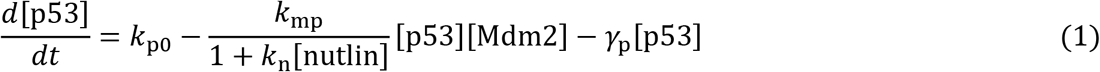

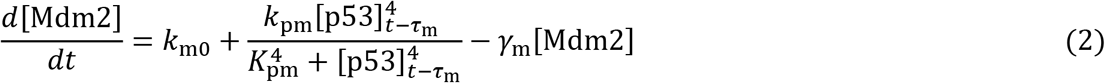

where [] denotes the dimensionless concentrations of the total proteins (p53 and Mdm2). *k*_p0_ (*k*_m0_) and *γ*_p_ (*γ*_m_) describe the rate of basal production and degradation of p53 (Mdm2), respectively. *k*_mp_ is the rate constant of Mdm2-mediated p53 degradation. The transcriptional activation of Mdm2 by the tetrameric p53 is characterized by a Hill function of 4^th^order with a rate constant *k*_pm_, Michaelis parameter *K*_pm_ and time delay *τ*_m_. [nutlin] is the concentration of Nutlin-3a in the unit of μM, and *k*_N_ is the inhibitory strength of Nutlin-3a in attenuating the Mdm2-mediated p53 degradation. We set *k*_N_ = 0.45 for all of the following simulation analysis, as it produced dose responses largely similar to the experimental data.

We first analyzed how p53 dynamics depended on the different kinetic parameters considered in Equations (1) and (2) by numerically simulating the DDEs in a large parameter space. This allowed us to pinpoint parameter sets that can generate periodic pulsing of p53 at 1 μM Nutlin-3a for further analysis. Under all such parameter sets, the simulated dose responses to increasing concentrations of Nutlin-3a produced similar p53 dynamics with a distinct mode of transitional dynamics, i.e., an initial p53 pulse followed by an elevated plateau (Fig. 2A). Such transitional dynamics agreed with what we experimentally observed in A549, A375 and 769-P cells under the 5-FU treatment, suggesting that the p53 responses induced by 5-FU in these three cell lines were largely mediated by the central p53-Mdm2 negative feedback loop. However, this mode of p53 transitional dynamics was not observed experimentally for the dose responses to Nutlin-3a or Etoposide. Moreover, we analyzed the distribution of the parameter values (Fig. 2B). The rate constants associated with p53 and Mdm2 interactions, including the p53-induced Mdm2 production rate (*k*_pm_) and the Mdm2-mediated p53 degradation rate (*k*_mp_) were most broadly distributed, spanning over a 30 to 50-fold range. This showed that the p53 pulsing output can be retained over a large variation of these two constants, indicating the interaction structure of the p53-Mdm2 negative feedback motif largely determines the pulsing output. Comparatively, the value for the Mdm2 basal degradation rates, *γ*_*m*_, exhibited the most narrow distribution, though still an 8-fold range. The output of the periodic pulsing mode of p53 is thsu likely most sensitive to cellular noise and/or cellular changes that alter this basal degradation rate.

**Figure 2.**
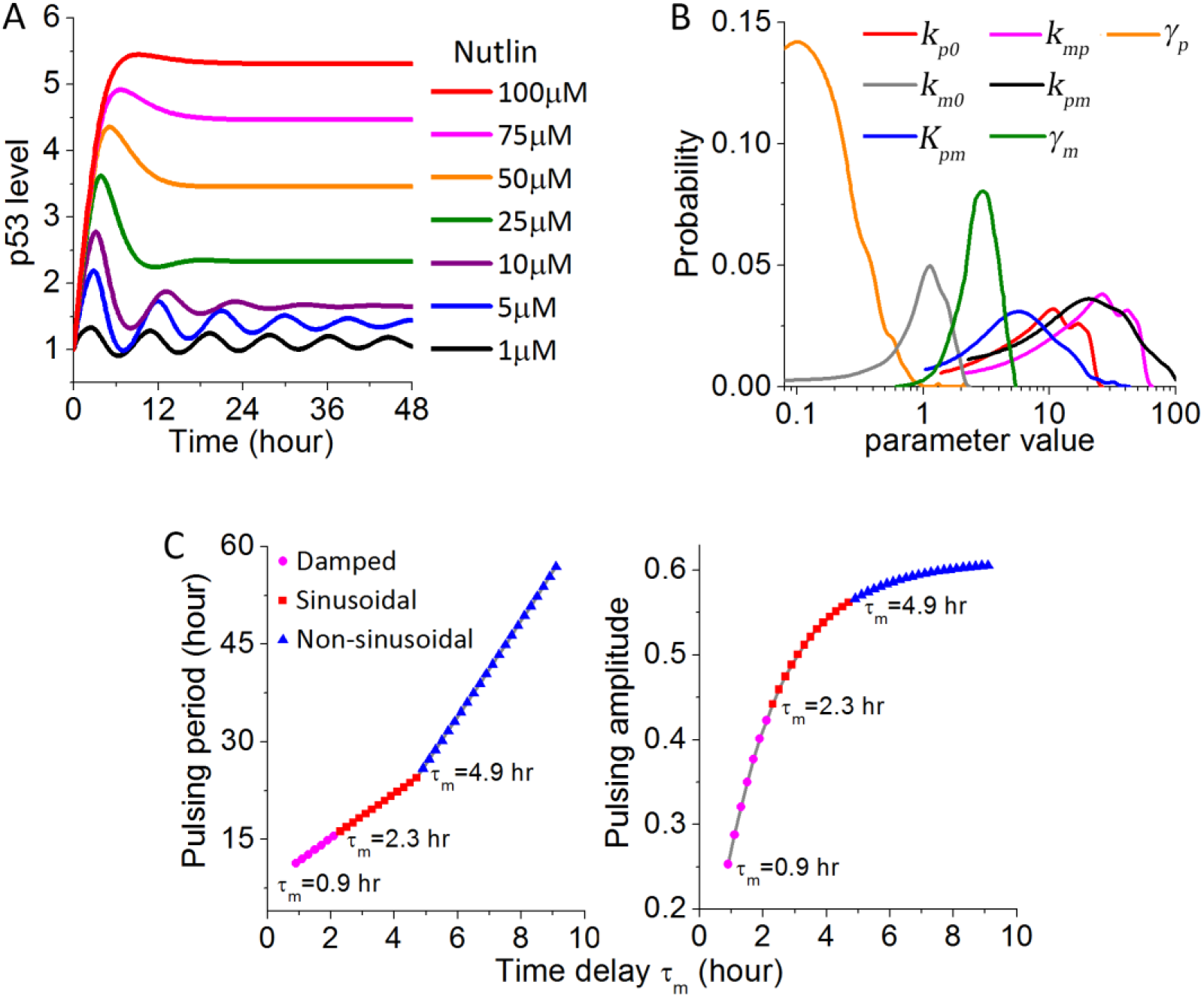
Modeling analysis of the dose-dependent outputs of the central p53-Mdm2 negative feedback showed a distinct mode of p53 transitional dynamics. (A) Simulation results of the dose response of p53 dynamics upon increasing Nutlin-3a concentrations. (B) Distributions of the values of the kinetic parameters involved in the p53-Mdm2 negative feedback as formulated in Equations (1) and (2), which can result in periodic pulsing of p53 at 1 μM Nutlin-3a. (C) Dependence of the p53 pulsing period (left panel) and pulsing amplitude (right panel) (at 1 μM Nutlin-3a) on the time delay in p53-induced Mdm2 upregulation, *τ*_m_,

Besides the above rate constants, dynamic output of the p53-Mdm2 negative feedback is also regulated by the time delay, *τ*_m_, in p53-mediated Mdm2 upregulation. In the above simulations, we set *τ*_m_ = 2.1 hr, similar to most previous modeling analysis. Varying the value of *τ*_m_ had a significant impact, in particular on the oscillatory features of p53 at low drug doses (Fig. 2C). Periodic oscillation of p53 was observed only when *τ*_m_ ≥ 0.9 hour (hr). For 0.9 hr ≤ *τ*_m_ < 2.3 hr, p53 dynamics displayed damped oscillation, while 2.3 hr ≤ *τ*_m_ < 4.9 hr gave rise to undamped, regular sinusoidal p53 oscillations. For *τ*_m_ ≥ 4.9 hr, p53 oscillation became non-sinusoidal, but still maintained the periodicity. Moreover, across the whole oscillatory regime, the pulsing period of p53 was found to be linearly proportional to the time delay, *τ*_m_, and the pulsing amplitude also showed positive dependence on the *τ*_m_ value (Fig. 2C). However, varying *τ*_m_ did not alter the transitional dynamics of p53 generated by the p53-Mdm2 negative feedback as shown in Figure 2A, again supporting that the interaction structure of the p53-Mdm2 negative feedback determines the dynamic phenotypes of p53 responses.

### p53 transitional dynamics conferred by additional positive feedback motifs

To identify the additional regulatory components and interaction motifs needed to generate the context-dependent p53 dynamics that we experimentally observed, we first examined the impact of additional positive feedback loops. For simplicity, we chose to analyze the following two positive feedback structures, where p53 transcriptionally activates a downstream target gene that positively enhances p53 upregulation either directly (Type 1) or by inhibiting Mdm2 (Type 2) (Fig. 3A). Here we showed simulation results of the dose-dependent p53 output in response to Nutlin-3a with these two different types of positive feedback motifs, as p53 dynamics induced by Nutlin-3a deviated most significantly from the simulated dynamic output of the p53-Mdm2 negative feedback alone. Kinetics of the positive feedback gene, denoted as PF and its direct activating activity on p53 or inhibitory activity on Mdm2 can be formulated as follows,

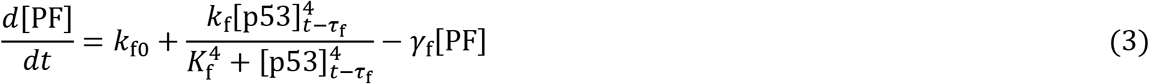

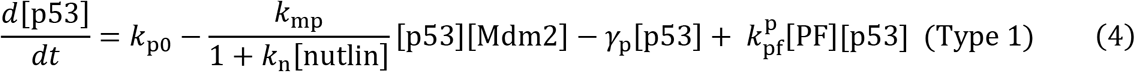

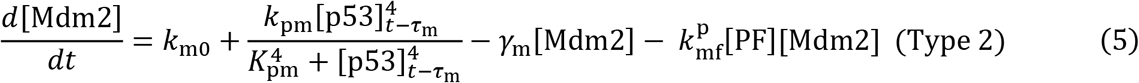

where *k*_f0_ and *γ*_f_ are the rate of basal production and degradation of the positive feedback gene, PF. The rate constant *k*_f_, Michaelis parameter *K*_f_ and time delay *τ*_f_ describe the transcriptional activation of PF by p53. 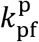 and 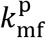 are the rate constant of p53-PF interaction in the Type 1 motif and the Mdm2-PF interaction in the Type 2 motif, respectively.

**Figure 3.**
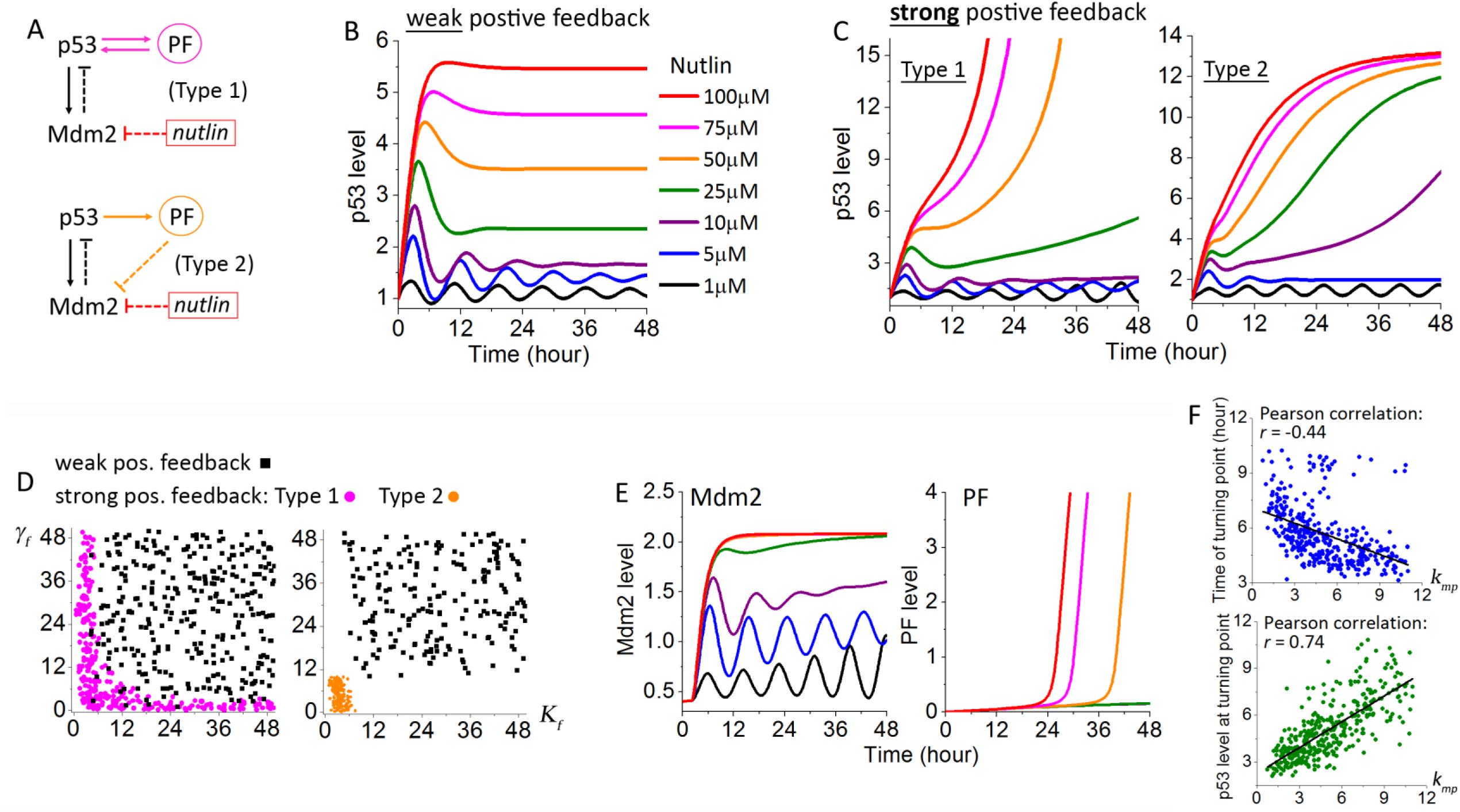
Strong positive feedback significantly alters the dose response of p53 dynamics, especially rendering unique p53 transitional dynamics. (A) Pathway diagrams with Type 1 or Type 2 positive feedback motif in addition to the central p53-Mdm2 negative feedback. PF denotes the positive feedback gene that is a transcriptional target of p53. (B, C) Dose response of p53 dynamics under (B) weak positive feedback strength and C) strong positive feedback strength (Left panel: Type 1; Right panel: Type 2). (D) Parameter space search showed the positive feedback strength is determined mainly by *γ*_f_ (the degradation rate of PF) and/or *K*_f_ (the Michaelis parameter for p53-induced PF upregulation). For Type 1 feedback motif (denoted in magenta), strong positive feedback can be achieved by either *γ*_f_ or *K*_f_ being small. For Type 2, both *γ*_f_ and *K*_f_ need to be small. (E) Dynamics of Mdm2 (left panel) and PF (right panel) resulted from the Type 1 positive feedback motif under increasing concentrations of Nutlin-3a. The drug concentrations are color coded as those in (B, C). (F) Correlation analysis of the time (upper panel) and p53 level (lower panel) of the turning point of p53 dynamics under intermediate concentrations of Nutlin-3a showed a significant dependence on *k*_mp_, the rate constant of Mdm2-mediated p53 degradation.

For both the Type 1 and Type 2 motifs, simulations over a large parameter space returned two distinct dose-response phenotypes of p53 dynamics. The first phenotype was common between the two motifs and largely similar to the output of the p53-Mdm2 negative feedback alone (Fig. 3B). Therefore we attributed it to a weak positive feedback strength. The second dynamic phenotype differed between the two motifs, though still sharing some common features. More importantly, they both diverged significantly from the p53-Mdm2 output, which we attributed to a strong positive feedback strength (Fig. 3C). For the Type 1 motif, the simulated p53 dynamics under strong positive feedback were largely similar to those that we experimentally observed in A549, U-2 OS and 769-P cells in response to Nutlin-3a. The novel p53 transitional dynamics at intermediate Nutlin-3a concentrations, i.e., an initial p53 pulse followed by an exponential increase of p53 level, were also reproduced. As for the Type 2 motif, it resulted in a different p53 transitional dynamics, in which the initial p53 pulse was followed by a sigmoidal increase of p53 level. This phenotype resembled the p53 response in U-2 OS cells under 5-FU. Interestingly, we found the positive feedback strength was mainly determined by the degradation rate of PF, *γ*_f_, and the Michaelis parameter, *K*_f_ (Fig. 3D). When either *γ*_f_ or *K*_f_ was small under the Type 1 motif, the second dose response phenotype of p53 dynamics (denoted as the “strong pos. feedback” phenotype) was produced from the model simulation. Under the Type 2 motif, the strong positive feedback phenotype required both *γ*_f_ and *K*_f_ to be small. This result makes quantitative sense, as the level of the positive feedback gene, PF, increase faster and higher when *γ*_f_ and/or *K*_f_ are small, thus resulting in stronger positive feedback effect on p53 induction.

The novel p53 transitional dynamics provide previously unexplored quantitative features to infer p53 pathway motifs and understand their functional impacts. Intuitively, the change of p53 dynamics from pulsing to an exponential or sigmoidal increase is due to the differential feedback contribution in time from the p53-Mdm2 negative feedback and the p53-PF positive. In the early phase of p53 upregulation, the inhibitory effect from Mdm2 dominates, restraining p53 in a low, pulsing mode (Fig. 3E). As Mdm2 levels to a saturating state, upregulation of the positive feedback gene, PF, continues. At the turning point, the activating effect from PF outweighs the inhibitory effect of Mdm2, thus causing p53 dynamics to change to a high-level induction mode. Correlation analysis of the turning point characteristics, including the time at which the p53 dynamic mode changes and the p53 level at the point of change, with the different rate constants showed that the turning point did not depend on the rate constants associated with PF (data not shown). This further indicated that the turning point is controlled by Mdm2. Specifically, we found the turning point depended most strongly on *k*_mp_, the rate constant of Mdm2-mediated p53 degradation, pointing to the inhibitory strength of Mdm2 over p53 as the determinant of the dynamic switch of p53 in time (Fig. 3F). Overall, our simulation analysis revealed a possible dynamic origin of the new p53 transitional dynamics that we observed in response to Nutlin-3a and 5-FU, and suggested that the therapeutic effects of these two drugs in some mammalian cell types are mediated by not only inhibiting Mdm2 but also activating additional positive feedback motif of the p53 pathway.

### p53 dynamics conferred by additional negative feedback motifs

Next we examined the quantitative impact of additional negative feedback motifs on top of the central p53-Mdm2 negative feedback loop. Again we considered two simple scenarios, where a p53 target gene, NF, attenuates p53 upregulation either directly (Type 1) or by activating Mdm2 (Type 2) (Fig. 4A). Mathematical formulation of the additional negative feedback is largely similar to Equations (3)–(5), except for changing the signs of the last term in Equations (4) and (5) to account for the inhibitory effect of NF on p53 level. As expected, simulations of these two negative feedback motifs under weak p53-NF or Mdm2-NF feedback strength produced the same p53 dynamics as discussed above for the p53-Mdm2 negative feedback alone as well as with the additional positive feedback motifs. Under strong NF negative feedback strength, both Type 1 and Type 2 motifs output an extended large pulse as drug concentrations increased, with Type 2 motif resulting in sharper pulsing features (Fig. 4B). This illustrated the general effect of negative feedback in p53 pathway to promote p53 pulsing, retraining overall p53 induction at a low level, which we previously found to promote cell cycle arrest over cell death [15]. Different from the positive feedback motifs, we found the negative feedback strength of the Type 2 motif was determined mainly by the basal production rate of NF (*k*_f0_) and the rate constant of Mdm2-NF interaction 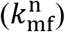, and both of these rate constants have to be large in order to render a strong negative feedback effect from the Type 2 motif (Fig. 4C).

**Figure 4.**
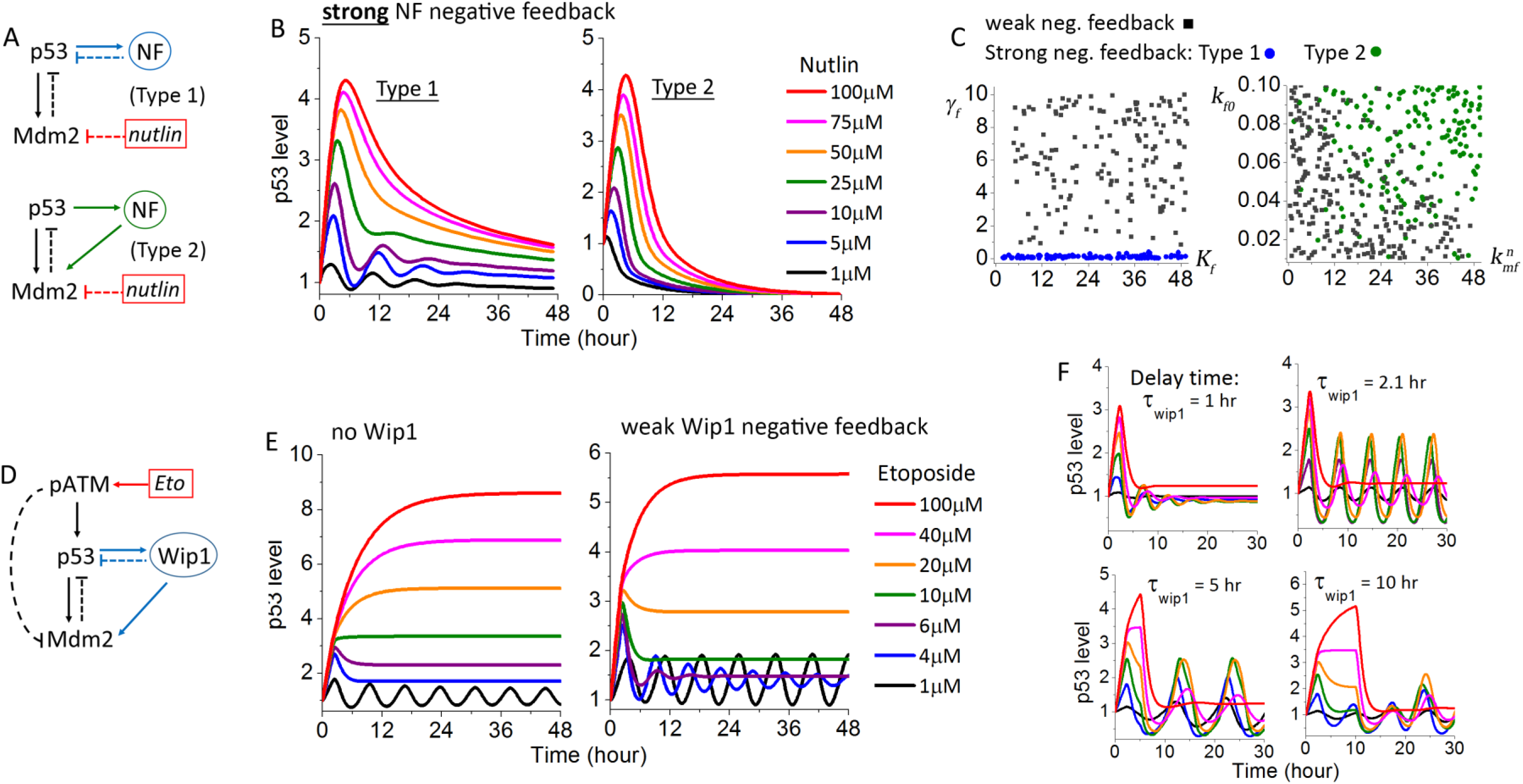
Additional negative feedbacks suppress p53 upregulation by enhancing the pulsing response of p53. (A) Pathway diagrams with Type 1 or Type 2 negative feedback motif in addition to the central p53-Mdm2 negative feedback. NF denotes the negative feedback gene that is a transcriptional target of p53. (B) Dose response of p53 dynamics under strong negative feedback strength (Left panel: Type 1; Right panel: Type 2). (C) The negative feedback strength is determined mainly by *γ*_f_ (the degradation rate of NF) for the Type 1 motif (denoted in blue), and *k*_f0_ (the basal production rate of NF) and 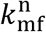 (the rate constant of Mdm2-NF interaction) for the Type 2 motif (green). (D). Pathway diagram of the ATM/p53/Mdm2/Wip1 regulatory module that mediates the cellular response to Etoposide. The phosphatase, Wip1, constitutes an additional, dual negative feedback of Type 1 plus Type 2 motifs. (E) Dose response of p53 dynamics in the absence of Wip1 (left panel) and in the presence of a weak Wip1 negative feedback (right panel). (F) The detailed features of p53 pulsing dynamics depend on the time delay in p53-induced Wip1 upregulation, *τ*_wip1_.

Of the five cancer cell lines that we profiled, MCF7 is evidently the one that strongly depends on the negative feedback interactions in the p53 pathway, as it exhibits p53 pulsing in response to all three drugs and even at high drug concentrations. In our previous study of the p53 dose response to Etoposide, we identified that the suppressive, pulsing p53 output seen in MCF7 was the result of a combination of downregulation in the ATM/p53/Mdm2 positive feed-forward activity and upregulation of the negative feedback via the phosphatase, Wip1 (Fig. 4D) [15]. As Wip1 dephosphorylates both p53 and Mdm2, it constitutes a dual negative feedback of Type 1 plus Type 2. Simulation results of the ATM/p53/Mdm2/Wip1 regulatory module under weak Wip1 negative feedback strength showed a p53 dose response with a more distinctive separation of the dynamic mode of p53 pulsing and monotonic induction. As shown in Figure 4E, the central ATM/p53/Mdm2 module without Wip1 generated p53 transitional dynamics in the form of an initial pulse followed by an elevated plateau. The presence of additional Wip1 negative feedback enhanced the pulsing characteristics at intermediate Etoposide concentrations, leading to a much lower steady-state plateau of p53 level. Given the optical and cellular noises in the experimental data, features of the p53 transitional dynamics may be further attenuated. Therefore, effects from additional weak negative feedbacks in the p53 pathway, such as via Wip1 or other negative feedback genes, may account for the bimodal p53 dynamics that we experimentally observed in the Etoposide response, i.e., the cell population shifted from p53 pulsing directly to monotonic induction without an observable different p53 transitional dynamics. Intuitively, a weak negative feedback strength can be the simple result of low expression level of the negative feedback gene, e.g., we noted that A549, U-2 OS and A375 all had low Wip1 expression [15].

In all simulation analysis described above, we set the time delay for the additional positive and negative feedback gene to be the same as that for Mdm2, i.e., *τ*_f_ = 2.1 hr. In general, we found decreasing the time delay *τ*_f_ in the positive feedback motifs accelerated the switching of p53 dynamics from pulsing to exponential/sigmoidal increase, while increasing the time delay (i.e., slowing down PF upregulation) delayed and attenuated the transition. As for the negative feedback motifs, shorter delay time further enhanced the suppressive effects of the negative feedback in attenuating both the p53 pulsing amplitude and the steady-state plateau level, e.g., similar to features experimentally observed in MCF7 cells (Fig. 4F, *τ*_wip1_ = 1 hr). A longer delay time prolonged the pulsing period and increased the pulsing amplitude, in particular for the first pulse and at high drug doses, which led to a more bimodal separation of the p53 dynamic mode of low periodic pulsing and an extended large pulse, similar to what we observed in 769-P cells under Etoposide (Fig. 4F, *τ*_wip1_ = 10 hr).

## Discussion

By integrating single cell dynamics data and mathematical modeling, our study not only characterized novel p53 dynamic modes and variable dose responses to distinct anticancer chemotherapeutics but also elucidated the underlying dynamic mechanisms for differential, context-dependent p53 dynamic control at the collective regulatory module level. We found unique p53 dynamics can be generated in a stimulus- and cell type-dependent manner by either positive or negative feedback motifs that co-opt with the central p53-Mdm2 negative feedback to modulate the signaling outputs. Although the feedback structures that we analyzed are simple, they are already able to explain a wide spectrum of p53 dose-response dynamics that we experimentally observed at the quantitative level. Our results suggest that taking a modular approach based on pathway interaction motifs is an effective way to dissect and investigate the dynamic outputs of complex signaling pathways. The pathway building blocks can also be employed to gain new mechanistic insight into drug mechanism and cell type variation that impacts drug sensitivity.

One important and reassuring finding from our modeling analysis is that the variable p53 dynamics are determined largely by the interaction structures, i.e., the types of feedback motifs involved and their wiring, and do not depend on the precise values of the interaction parameters. For instance, the distinct p53 dynamic modes activated by the two types of positive feedback motifs at intermediate drug concentrations are invariant over a large parameter space, as long as the feedback strength is strong. The confirmation that the interaction structure determines the dynamic output of signaling pathways provides further support of the robustness of biological networks. That is, signaling response of large networks/pathways and their functional consequences can remain relatively stable, despite the presence of constant cellular noises and small environmental fluctuations that likely alter the values of multiple pathway parameters. Our results also suggest that the specific choice of parameter values in the modeling analysis is unlikely to affect the theoretical results in terms of the differential dynamic phenotypes of p53, although the detailed quantitative features, e.g., pulsing period and amplitude, may vary. In a literature search, we found parameter values for a few key p53 pathway interactions used in different modeling papers can span two orders of magnitude. Our work provides a quantitative explanation why similar p53 dynamics can be generated with such largely different parameter values as well as a basis to compare the modeling results from these previous studies.

The cell-type variation in p53 dynamic response presents a new angle to identify the points of variation in different cancer cell types that confer differential drug sensitivity. Tumors are known to be highly heterogeneous both at the genotype and phenotype level. Efforts to connect genotype with phenotype so far are not successful partly because dynamic output of the signaling networks, as we showed here, can be altered by collective, moderate changes in multiple network components, rather than a large gain- or loss-of-function mutation in a single gene. We believe using the modular approach that profiles the variable dependence of different tumor subtypes on key interaction motifs could be an alternative way to bioinformatics analysis of large network to uncover the mechanistic origins of phenotypic heterogeneity in cancer. Moreover, dose dependent dynamics of key signaling proteins, such as p53, can be a convenient readout to probe the cell-type dependent modular output and identify the underlying points of variation.

The fact that p53 dynamics induced by Nutlin-3a cannot be recapitulated by only inhibiting Mdm2 in the p53-Mdm2 negative feedback revealed that the drug action of Nutlin-3a involves additional p53 pathway component(s) that are yet to be elucidated. We showed that in three out of the five cell lines, Nutlin-3a upregulates p53 by activating a direct positive feedback (Type 1) to enhance p53 induction in addition to inhibiting Mdm2. However, it is unclear what p53 pathway component constitutes this Type 1 positive feedback to mediate the Nutlin-3a response. Most reported p53 positive feedback motifs are of the Type 2 configuration, such as the positive feedbacks conferred by p21, PTEN and miRNAs [28–30]. We nonetheless attempted to experimentally examine the involvement of PTEN, p21 and miR605 in the response of A549 to Nutlin-3a. Preliminary data did not identify a key regulatory role from these positive feedback components to mediate Nutlin-3a activity. The molecular identity of the additional Type 1 positive feedback component involved in the Nutlin-3a response is still yet to be determined in further study. The p53 dynamics and cell lines that we identified to exhibit the novel p53 transitional dynamics under Nutlin-3a can provide the model system to explore this intriguing Type 1 positive feedback target gene of p53.

## Materials and Methods

### Cell culture and Chemicals

Cell lines were purchased from American Type Culture Collection (ATCC, USA) and cultured under 37°C and 5% CO_2_ in appropriate medium supplemented with 10% Fetal Calf Serum (FCS), 100U/ml penicillin and 100μg/ml streptomycin. A549 was maintained in F-12K, A375 in DMEM, U-2 OS in McCoy’s, MCF7 and 769-P in RPMI. To generate fluorescent reporter cells for live-cell imaging of real-time p53 dynamics, we infected each cell line with lentiviruses encoding an established p53-Venus reporter and selected isogenic clones that exhibited drug response most similar to their respective parental line for conducting the single cell imaging experiments. The p53-venus reporter, consisting of wild-type p53 fused to a yellow fluorescent protein, Venus, was a generous gift from Dr. Galit Lahav (Department of Systems Biology, Harvard Medical School). Etoposide, Nutlin-3a and 5-Fluouracil were purchased from Tocris.

### Time-lapse microscopy

Cells were plated in 24-well imaging dish (Cellvis, USA) and cultured in phenol red-free CO_2_-independent medium (Invitrogen) supplemented with 10% FCS, 100 U/ml penicillin and 100 μg/ml streptomycin. Cell images were acquired with the Nikon Ti2-E inverted microscope enclosed in a humidified chamber maintained at 37°C. Cells were imaged every 10 minutes using a motorized stage and a 20X objective (NA=0.95).

The real-time dynamics of nuclear p53 was scored based on the p53-Venus fluorescence in the nucleus. For quantifying the single-cell p53 traces, we used an automatic cell tracking program that we developed using Matlab. The program consists of image analysis procedures that sequentially segment the individual cells, track them in time, identify the nucleus and measure the p53 fluorescence intensity in the nucleus.

### Mathematical models and computational analysis

Detailed description of the mathematical models, computational simulations and parameter sets of the models can be found in the result section and the supplementary materials.

## Author contributions

J.S. designed and supervised the study; J.X. developed the mathematical model and conducted the computational analysis; L.Z., Y.Z. and X.L. performed the experiments; J.X. and J.S. analyzed the data; J.X. and J.S. wrote the paper.

## Acknowledgements

We thank Dr. Galit Lahav (Department of Systems Biology, Harvard Medical School) for the p53-Venus lenti-viral vector. This work was supported by the Hong Kong Research Grant Council (#N_HKBU215/13, #T12-710/16-R and #C2006-17E) to J. Shi and the “Laboratory for Synthetic Chemistry and Chemical Biology” under the Health@InnoHK Program by the Innovation and Technology Commission of Hong Kong. The authors declare no conflict of interest.

