## Supplementary materials for "Single-cell transitional dynamics unravel stimulus- and cell type-dependent signaling outputs of distinct p53 regulatory feedback motifs"

**This file includes:**

Supplementary text describing the mathematical model, simulation and parameters.

Table S1. Model parameters for simulating the p53 dynamic responses to Nutlin-3a.

Table S2. Model parameters for the additional positive feedback motifs associated with PF.

Table S3. Model parameters for the additional negative feedback motifs associated with NF.

Table S4. Model parameters for simulating the p53 dynamic output of the ATM/p53/Mdm2/Wip1 regulatory module in response to Etoposide.

Figure S1. Simulation results for the ATM/p53/Mdm2 module in response to Etoposide.

### I. Mathematical model and analysis of the ATM/p53/Mdm2 regulatory module in response to Etoposide

Dynamic output of the p53-Mdm2 motif in response to Etoposide is formulated by the following simplified delay differential equations (DDEs).

$$\frac{d[\mathrm{ATM_p}]}{dt} = S_{\mathrm{ea}}[\mathrm{Eto}]\left([\mathrm{ATM_t}] - [\mathrm{ATM_p}]\right) - D_{\mathrm{a0}}[\mathrm{ATM_p}] \tag{1}$$

$$\frac{d[\mathrm{p53}]}{dt} = k_{\mathrm{p0}} - k_{\mathrm{mp}}\left([\mathrm{ATM_p}]\right)[\mathrm{p53}][\mathrm{Mdm2}] - \gamma_{\mathrm{p}}[\mathrm{p53}] \tag{2}$$

$$\frac{d[\mathrm{Mdm2}]}{dt} = k_{\mathrm{m0}} + \frac{k_{\mathrm{pm}}[\mathrm{p53}]_{t-\tau_{\mathrm{m}}}^{4}}{K_{\mathrm{pm}}^{4}\left([\mathrm{ATM_p}]_{t-\tau_{\mathrm{m}}}\right) + [\mathrm{p53}]_{t-\tau_{\mathrm{m}}}^{4}} - \gamma_{\mathrm{m}}\left([\mathrm{ATM_p}]\right)[\mathrm{Mdm2}] \tag{3}$$

where [ ] denotes dimensionless concentrations of the total proteins (p53, Mdm2, $\mathrm{ATM_t}$) or the active, phosphorylated form of ATM ($\mathrm{ATM_p}$). Briefly, ATM is phosphorylated and activated by different concentrations of Etoposide with rate constant $S_{\mathrm{ea}}$ and $\mathrm{ATM_p}$ is dephosphorylated with rate constant $D_{\mathrm{a0}}$ (Equation (1)). $\mathrm{ATM_p}$ subsequently phosphorylates p53 and Mdm2, leading to decrease in p53-Mdm2 binding and Mdm2-mediated p53 degradation, as described by the $\mathrm{ATM_p}$-dependent rate parameter $k_{\mathrm{mp}}$. $k_{\mathrm{p0}}$ and $\gamma_{\mathrm{p}}$ describe the rate of basal production and Mdm2-independent degradation of p53, respectively (Equation (2)). The transcriptional activation of Mdm2 by tetrameric p53 is characterized by a Hill function of $4^{\mathrm{th}}$ order in Equation (3) with rate constant $k_{\mathrm{pm}}$, Michaelis parameter $K_{\mathrm{pm}}$ and time delay $\tau_{\mathrm{m}}$. Our previous work showed that $\mathrm{ATM_p}$ enhanced auto-degradation of the phosphorylated form of Mdm2, therefore the Mdm2 degradation rate $\gamma_m$ is also set to be $\mathrm{ATM_p}$ dependent [15].

Equations (1)-(3) were derived with the assumption of rapid equilibrium of phosphorylation and dephosphorylation, as they are much faster than transcription and degradation.

This assumption allowed us to simplify the kinetic equations for total p53 and Mdm2 without specifying their respective phosphorylated and unphosphorylated forms, as well as obtain the $ATM_p$ dependence of the kinetic rates as follows (refer to the supplementary information of ref. [15] for the detailed derivation).

$$k_{\mathrm{mp}}([\mathrm{ATM_p}]) = k_{\mathrm{mp0}}(\frac{D_{\mathrm{p0}}}{S_{\mathrm{p0}} + D_{\mathrm{p0}} + S_{\mathrm{ap}}[\mathrm{ATM_p}]}) \quad (4)$$

$$K_{\mathrm{pm}}([\mathrm{ATM_p}]) = K_{\mathrm{pm0}}(1 + \frac{D_{\mathrm{p0}}}{S_{\mathrm{p0}} + S_{\mathrm{ap}}[\mathrm{ATM_p}]}) \quad (5)$$

$$\gamma_{\mathrm{m}}([\mathrm{ATM_p}]) = \frac{\gamma_{\mathrm{m0}} D_{\mathrm{m0}} + \gamma_{\mathrm{m1}}(S_{\mathrm{m0}} + S_{\mathrm{am}}[\mathrm{ATM_p}])}{D_{\mathrm{m0}} + S_{\mathrm{m0}} + +S_{\mathrm{am}}[\mathrm{ATM_p}]} \quad (6)$$

where $S_{\mathrm{p0}}$ ($S_{m0}$) and $D_{\mathrm{p0}}$ ($D_{m0}$) are the basal phosphorylation and dephosphorylation rate constants of p53 (Mdm2), $S_{\mathrm{ap}}$ ($S_{am}$) are the ATM-mediated phosphorylation rate of p53 (Mdm2), $k_{\mathrm{mp0}}$ is the Mdm2-mediated p53 degradation rate constant, $K_{\mathrm{pm0}}$ is the Michaelis constant for p53-induced upregulation of Mdm2, and $\gamma_{m0}$ and $\gamma_{m1}$ are the degradation rate constants of the unphosphorylated and phosohorylated Mdm2, respectively.

Similar to the modeling results for the p53-Mdm2 negative feedback motif under Nutlin-3a, p53 levels simulated for the ATM/p53/Mdm2 module produced a unique p53 transitional dynamics at intermediate Etoposide concentration, i.e., an initial large p53 pulse followed by an elevated plateau (supplementary Fig. S1A). This again did not agree with our experimental data for the Etoposide response and suggested additional regulatory components/interactions are involved beyond the ATM/p53/Mdm2 interactions. Figure S1B showed the distribution of parameter values that can generate periodic pulsing of p53 at 1 μM Etoposide as well as monotonic p53 induction at high drug dose, as we observed experimentally for the etoposide-sensitive cell lines. The values for rate constants of Mdm2-mediated p53 degradation ($k_{\mathrm{mp0}}$), p53-induced Mdm2 production ($k_{\mathrm{pm}}$) and ATM-mediated Mdm2 degradation ($\gamma_{m1}$) were mostly broadly distributed, while the Michaelis constant $K_{\mathrm{pm0}}$ spanned a smaller, but still 10-fold range.

As expected, the dynamic output of ATM/p53/Mdm2 is also regulated by the time delay in p53-mediated Mdm2 upregulation, $\tau_{\mathrm{m}}$. By varying $\tau_{\mathrm{m}}$, we found the oscillatory feature of p53 level at low drug dose is evident only when $\tau_{\mathrm{m}} \geq$ 0.7 hour (hr) (Fig. S1C, left panel). More specifically, when 0.7 hr $\leq \tau_{\mathrm{m}} <$ 1.7 hr, p53 dynamics exhibit damped oscillation, while 1.7 hr $\leq \tau_{\mathrm{m}} <$ 4.3 hr gives rise to steady sinusoidal p53 oscillation. For $\tau_{\mathrm{m}} \geq$ 4.3 hr, p53 oscillation becomes non-sinusoidal, but maintains the periodicity. Across the whole oscillatory regime, period of the p53 oscillation is proportional to the time delay $\tau_{\mathrm{m}}$, and the oscillation amplitude also showed largely linear proportionality with $\tau_{\mathrm{m}}$, when $\tau_{\mathrm{m}} <$ 4.3 hr (Fig. S1C, right panel). However, varying $\tau_{\mathrm{m}}$ again did not alter the transition dynamics of p53 induction as discussed above.

### II. Parameters and simulation

For parameter space search, we randomly generated large sets of parameter values and then selected those that can result in periodic pulsing of p53 at low drug dose for further analysis. The representative p53 dynamics shown in the figures were acquired by simulating the respective mathematical models with the parameter sets shown in the supplementary Table S1-S4. These representative parameters were chosen based on our imaging data and western blot data in a previous study [15]. And we set the parameters such that the system stays in a stable state at [Drug] = 0 and time = 0, with [p53]=1, [Mdm2]=0.4, [PF]=0 or [NF]=0, [$ATM_p$]=0 and [Wip1]=0.3. The above initial values of protein concentrations were employed for all simulations. The model simulations were performed using the Matlab built-in function dde23.

**Supplementary Table S1: Model parameters for simulating the p53 responses induced by Nutlin-3a**

| Parameter | Value & unit | Interpretation |
|---|---|---|
| $k_{p0}$ | 1.3396 $h^{-1}$ | Basal production rate of p53 |
| $k_{mp0}$ | 3.0285 $h^{-1}$ | Rate constant of Mdm2-induced degradation of p53 |
| $\gamma_p$ | 0.1 $h^{-1}$ | Mdm2-independent degradation rate constant of p53 |
| $k_{m0}$ | 0.08 $h^{-1}$ | Basal production rate of Mdm2 |
| $k_{pm}$ | 0.9172 $h^{-1}$ | Rate constant of p53-induced production of Mdm2 |
| $K_{pm0}$ | 1.61 | Michaelis constant for p53-induced production of Mdm2 |
| $\gamma_m$ | 0.4799 $h^{-1}$ | Degradation rate constant of Mdm2 |
| $\tau_m$ | 2.1 h | Time delay in production of Mdm2 |

**Supplementary Table S2: Model parameters for the additional positive feedback motifs associated with PF (data shown in Figure 3C and 3E)**

**Type 1 motif:**

| Parameter | Value & unit | Interpretation |
|---|---|---|
| $k_{f0}$ | 0.005 $h^{-1}$ | Basal production rate of PF |
| $k_f$ | 1 $h^{-1}$ | Rate constant of p53-induced production of PF |
| $K_f$ | 10.74 | Michaelis constant for p53-induced production of PF |
| $\gamma_f$ | 0.02 $h^{-1}$ | Degradation rate constant of PF |
| $k_{pf}^{p}$ | 11.5 $h^{-1}$ | Rate constant of p53 enhancement by PF |
| $\tau_f$ | 2.1 h | Time delay in production of PF |

**Type 2 motif:**

| Parameter | Value & unit | Interpretation |
|---|---|---|
| $k_{f0}$ | 0.07 $h^{-1}$ | Basal production rate of PF |
| $k_f$ | 12.16 $h^{-1}$ | Rate constant of p53-induced production of PF |
| $K_f$ | 2.4567 | Michaelis constant for p53-induced production of PF |
| $\gamma_f$ | 0.2462 $h^{-1}$ | Degradation rate constant of PF |
| $k_{mf}^{p}$ | 2.1347 $h^{-1}$ | Rate constant of Mdm2 inhibition by PF |
| $\tau_f$ | 2.1 h | Time delay in production of PF |

**Supplementary table S3: Model parameters for the additional negative feedback motifs associated with NF (data shown in Figure 4B)**

**Type 1 motif:**

| Parameter | Value & unit | Interpretation |
|---|---|---|
| $\boldsymbol{k}_{\mathbf{f0}}$ | 0.026 $h^{-1}$ | Basal production rate of NF |
| $\boldsymbol{k}_{\mathbf{f}}$ | 0.2831 $h^{-1}$ | Rate constant of p53-induced production of NF |
| $\boldsymbol{K}_{\mathbf{f}}$ | 39.448 | Michaelis constant for p53-induced production of NF |
| $\boldsymbol{\gamma}_{\mathbf{f}}$ | 0.018 $h^{-1}$ | Degradation rate constant of NF |
| $\boldsymbol{k}_{\mathbf{pf}}^{\mathbf{n}}$ | 3.096 $h^{-1}$ | Rate constant of p53 inhibition by NF |
| $\boldsymbol{\tau}_{\mathbf{f}}$ | 2.1 h | Time delay in production of NF |

**Type 2 motif:**

| Parameter | Value & unit | Interpretation |
|---|---|---|
| $\boldsymbol{k}_{\mathbf{f0}}$ | 0.094 $h^{-1}$ | Basal production rate of NF |
| $\boldsymbol{k}_{\mathbf{f}}$ | 36.14 $h^{-1}$ | Rate constant of p53-induced production of NF |
| $\boldsymbol{K}_{\mathbf{f}}$ | 22.646 | Michaelis constant for p53-induced production of NF |
| $\boldsymbol{\gamma}_{\mathbf{f}}$ | 3.171 $h^{-1}$ | Degradation rate constant of NF |
| $\boldsymbol{k}_{\mathbf{mf}}^{\mathbf{n}}$ | 19.83 $h^{-1}$ | Rate constant of Mdm2 enhancement by NF |
| $\boldsymbol{\tau}_{\mathbf{f}}$ | 2.1 h | Time delay in production of NF |

**Supplementary table S4: Model parameters for simulating the p53 dynamic output of the ATM/p53/Mdm2/Wip1 regulatory module in response to Etoposide**

| Parameter | Value & unit | Interpretation |
|---|---|---|
| $k_{p0}$ | 1.3396 $h^{-1}$ | Basal production rate of p53 |
| $k_{m0}$ | 0.08 $h^{-1}$ | Basal production rate of Mdm2 |
| $k_{pm}$ | 0.9172 $h^{-1}$ | Rate constant of $p53_p$-induced production of Mdm2 |
| $K_{pm0}$ | 0.4025 | Michaelis constant for $p53_p$-induced production of Mdm2 |
| $\tau_m$ | 2.1 h | Time delay in production of Mdm2 |
| $k_{mp0}$ | 4.038 $h^{-1}$ | Rate constant of Mdm2-induced degradation of $p53_u$ |
| $\gamma_p$ | 0.1 $h^{-1}$ | Mdm2-independent degradation rate constant of p53 |
| $\gamma_{m0}$ | 0.2579 $h^{-1}$ | Degradation rate constant of $Mdm2_u$ |
| $\gamma_{m1}$ | 7.7362 $h^{-1}$ | Degradation rate constant of $Mdm2_p$ |
| $S_{ea}$ | 1 $(\mu M \cdot h)^{-1}$ | Rate constant of etoposide-induced phosphorylation of ATM |
| $D_{a0}$ | 80 $h^{-1}$ | Rate constant of dephosphorylation of ATM |
| $S_{p0}$ | 0.25 $h^{-1}$ | Rate constant of ATM-independent phosphorylation of p53 |
| $S_{ap}$ | 0.8335 $h^{-1}$ | Rate constant of ATM-mediated phosphorylation of p53 |
| $S_{m0}$ | 0.0153 $h^{-1}$ | Rate constant of ATM-independent phosphorylation of Mdm2 |
| $S_{am}$ | 0.283 $h^{-1}$ | Rate constant of ATM-mediated phosphorylation of Mdm2 |
| $ATM_t$ | 16 | Total amount of ATM |

**Weak Wip1 negative feedback strength (data shown Figure 4E)**

| Parameter | Value & unit | Interpretation |
|---|---|---|
| $k_{w0}$ | 7.267 $h^{-1}$ | Basal production rate of Wip1 |
| $k_{pw}$ | 1.9067 $h^{-1}$ | Rate constant of $p53_p$-induced production of Wip1 |
| $K_{pw0}$ | 0.4025 | Michaelis constant for $p53_p$-induced production of Wip1 |
| $\gamma_w$ | 13.964 $h^{-1}$ | Degradation rate constant of Wip1 |
| $D_{p0}$ | 0.75 $h^{-1}$ | Rate constant of Wip1-independent dephosphorylation of p53 |
| $D_{wp}$ | 18.093 $h^{-1}$ | Rate constant of Wip1-mediated dephosphorylation of p53 |
| $D_{m0}$ | 0.5 $h^{-1}$ | Rate constant of Wip1-independent dephosphorylation of Mdm2 |
| $D_{wm}$ | 30.525 $h^{-1}$ | Rate constant of Wip1-mediated dephosphorylation of Mdm2 |
| $\tau_w$ | 2.1 h | Time delay in production of Mdm2 |

**Strong Wip1 negative feedback strength (data shown in Figure 4F)**

| Parameter | Value & unit | Interpretation |
|---|---|---|
| $\boldsymbol{k}_{\mathbf{w0}}$ | 2.377 $h^{-1}$ | Basal production rate of Wip1 |
| $\boldsymbol{k}_{\mathbf{pw}}$ | 24.932 $h^{-1}$ | Rate constant of $p53_p$-induced production of Wip1 |
| $\boldsymbol{K}_{\mathbf{pw0}}$ | 0.4025 | Michaelis constant for $p53_p$-induced production of Wip1 |
| $\boldsymbol{\gamma}_{\mathbf{w}}$ | 5.964 $h^{-1}$ | Degradation rate constant of Wip1 |
| $\boldsymbol{D}_{\mathbf{p0}}$ | 0.75 $h^{-1}$ | Rate constant of Wip1-independent dephosphorylation of p53 |
| $\boldsymbol{D}_{\mathbf{wp}}$ | 7.7107 $h^{-1}$ | Rate constant of Wip1-mediated dephosphorylation of p53 |
| $\boldsymbol{D}_{\mathbf{m0}}$ | 0.5 $h^{-1}$ | Rate constant of Wip1-independent dephosphorylation of Mdm2 |
| $\boldsymbol{D}_{\mathbf{wm}}$ | 33.118 $h^{-1}$ | Rate constant of Wip1-mediated dephosphorylation of Mdm2 |

### III. Supplementary figure

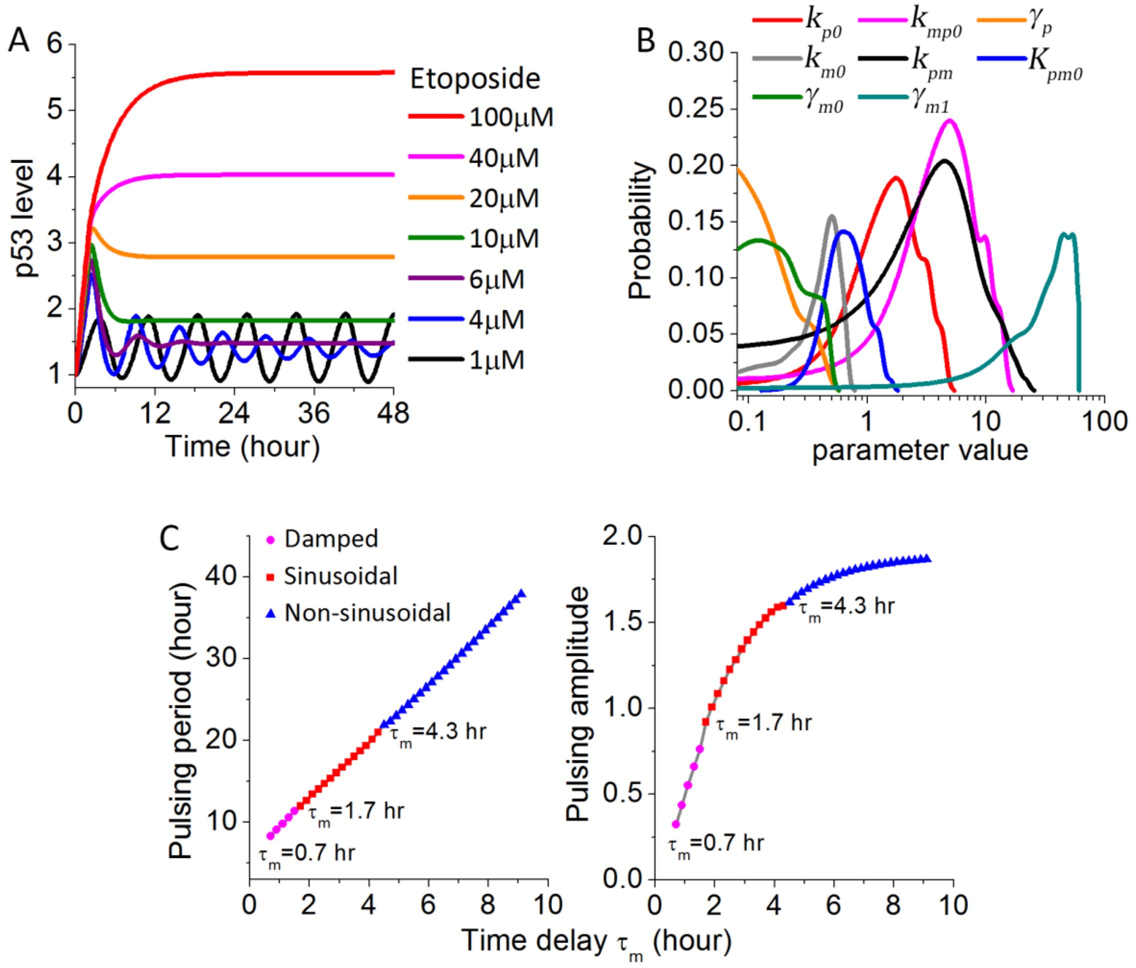


**Supplementary Figure S1.** Simulation results for the ATM/p53/Mdm2 module in response to Etoposide. (A) Simulation results of the dose response of p53 dynamics upon increasing Etoposide concentrations. (B) Distributions of the values of the kinetic parameters involved in the ATM/p53/Mdm2 regulatory module as formulated in Equations (1)-(3) in the supplementary text, which can result in periodic pulsing of p53 at 1 μM Etoposide. (C) Dependence of the p53 pulsing period (left panel) and pulsing amplitude (right panel) on the time delay in p53-mediated Mdm2 upregulation, $\tau_{\mathrm{m}}$, at 1 μM Etoposide.
